# Expanding the *BonnMu* sequence-indexed repository of transposon induced maize (*Zea mays* L.) mutations in dent and flint germplasm

**DOI:** 10.1101/2024.02.24.581857

**Authors:** Yan Naing Win, Tyll Stöcker, Xuelian Du, Alexa Brox, Marion Pitz, Alina Klaus, Heiko Schoof, Frank Hochholdinger, Caroline Marcon

## Abstract

The *BonnMu* resource is a transposon-tagged mutant collection designed for functional genomics studies in maize. To expand this resource, we crossed an active *Mutator* (*Mu*) line with dent (B73, Co125) and flint (DK105, EP1 and F7) germplasm, resulting in the generation of 8,064 mutagenized *BonnMu* F_2_-families. Sequencing of these *Mu*-tagged families revealed 425,924 heritable *Mu* insertions affecting 36,612 (83%) of the 44,303 high-confidence gene models of maize (B73v5). On average, we observed 12 *Mu* insertions per gene (425,924 total insertions/ 36,612 affected genes) and 53 insertions per *BonnMu* F_2_-family (425,924 total insertions/ 8,064 families). *Mu* insertions and photos of seedling phenotypes from segregating *BonnMu* F_2_-families can be accessed through the Maize Genetics and Genomics Database (MaizeGDB). Downstream examination via the automated Mutant-seq Workflow Utility (MuWU) identified 94% of the germinal insertion sites in genic regions and only a small fraction of 6% inserting in non-coding intergenic sequences of the genome. Consistently, *Mu* insertions aligned with gene-dense chromosomal arms. In total, 42% of all *BonnMu* insertions were located in the 5’ untranslated region (UTR) of genes, corresponding to accessible chromatin. Furthermore, for 38% of the insertions (163,843 of 425,924 total insertions) *Mu1*, *Mu8* and *MuDR* were confirmed to be the causal *Mu* elements. Our publicly accessible European *BonnMu* resource has archived insertions covering two major germplasm groups, thus facilitating both forward and reverse genetics studies.

## Introduction

Maize (*Zea mays* L.) has a long history of genetic investigation. Since the early 1900’s geneticists are collecting and describing maize mutations affecting a broad range of biological processes. However, to date, only a few hundred genes have been functionally characterized based on visible morphological mutant phenotypes (Schnable and Freeling, 2011; https://www.maizegdb.org). Forward genetic experiments represent a classical method to unravel gene functions by cloning a gene based on a mutant phenotype. In contrast, reverse genetic screens enable the identification of mutant phenotypes by utilizing disrupted gene sequences (Candela and Hake, 2008). Both approaches have been successfully applied in the past exemplified by their application in identifying and characterizing multiple genes underlying maize root development (Nestler *et al*., 2014; Xu *et al*., 2015; Li *et al*., 2016).

Genome-wide insertional mutagenesis represents a powerful reverse genetics tool to generate loss-of-function mutations for virtually all genes within a genome. Meanwhile, several maize genomes of two major germplasm, such as B73 and W22 of the dent pool (Schnable *et al*., 2009; Jiao *et al*., 2017; Springer *et al*., 2018) and F7, DK105 and EP1 of the European flint pool (Haberer *et al*., 2020) were sequenced. Dent and flint pools are genetically divergent with respect to several traits including early vigor and cold tolerance, which is due to their historic geographical separation and adaptation to different environmental conditions. The progenitors of the European flint pool reached higher latitudes, which required selection for cold tolerance and early maturation (Haberer *et al*., 2020). Therefore, the flint lines are important genetic resources for central European maize research due to their stable growth properties under temperate climatic conditions.

To facilitate genome-wide insertional mutagenesis screens in maize, *Mutator* (*Mu*) transposons are used as biological mutagens, because they can move from one location in the genome to another, thereby disrupting genes (Lisch, 2015). Transposons, also known as transposable elements, are mobile DNA sequences first discovered in maize (McClintock, 1951). They are classified in two major classes: class I retrotransposons, which necessitate an RNA intermediate for transposition and class II DNA transposons, which directly transpose via DNA transposase. *Mu* transposons, the most active class II transposon family in maize, consist of an autonomous element (*MuDR*; Robertson, 1978) and multiple non-autonomous elements (Lisch, 2002, 2015; Tan *et al*., 2011). All identified *Mu* transposons conserve highly similar 215 bp terminal inverted repeats (TIRs) at both ends of the elements and create 9 bp target site duplications directly flanking the *Mu* transposon sequences upon insertion. *Mu* elements randomly target genes throughout the maize genome (Lisch, 2015). As such, *Mu* insertion site frequencies were observed to strongly correlate with gene density (Schnable *et al*., 2009). To date, three public sequence-indexed mutant libraries have been established by *Mu* transposon insertional mutagenesis as invaluable resources for conducting functional genetics studies in maize (*UniformMu*: McCarty *et al*., 2013; *ChinaMu*: Liang *et al*., 2019; *BonnMu*: Marcon *et al*., 2020). These mutant collections are ideal starting points for forward and reverse genetic screens (1) to functionally characterize novel mutants regulating various developmental processes (Hunter *et al*., 2014; Dai *et al*., 2021) and (2) to validate candidate genes by additional allelic mutations. The *BonnMu* resource utilizes random *Mu* insertions to disrupt genes, based on a method called Mutant-seq (Mu-seq; McCarty *et al*., 2013). Mu-seq enables the identification of maize F_2_-families carrying transposon insertions in a sequence-indexed (i.e. transposon tagged) population, by high-throughput next-generation sequencing. The analysis of this European-based sequence-indexed resource has been optimized and accelerated by the MuWU bioinformatic pipeline (Stöcker *et al*., 2022), which facilitates unbiased and high-throughput sequencing of mutagenized *BonnMu* F_2_*-*families.

In this study, we extend the original *BonnMu* reverse genetics resource introduced by Marcon *et al*. in 2020. We achieved this expansion by sequencing an additional 6,912 mutagenized F_2_-families across various genetic backgrounds, including B73, Co125, DK105, EP1 and F7. Utilizing a consistent downstream analysis of all Mu-seq reads through the MuWU bioinformatic pipeline, we identified transposon-induced mutations in 83% of all maize genes. The enhanced *BonnMu* resource is now available for maize geneticists, providing an invaluable tool for molecular and genetic analyses.

## Results

### *BonnMu* insertions cover 83% of all annotated B73v5 gene models

The *BonnMu* F_2_-families were generated in different genetic germplasm backgrounds, i.e., B73 and Co125 from the dent pool and F7, EP1 and DK105 from the flint pool (Table S1). In a previous study, we generated two Mu-seq libraries comprising 1,152 *BonnMu* F_2_-families in B73 background (Marcon *et al*., 2020). For the downstream bioinformatic analysis, we employed the Mu-seq method described by McCarty *et al*., (2013) and Liu *et al*., (2016). In the present study, we complement the two previous Mu-seq libraries by 12 additional libraries comprising 6,912 *BonnMu* F_2_-families. To ensure equal analysis of all 14 datasets, we applied the bioinformatics method MuWU (Stöcker *et al*., 2022) and integrated the first two libraries (Marcon *et al*., 2020) into the analysis. Sequencing of 14 Mu-seq libraries yielded 1,589,204,597 raw read pairs (Table 1). After automated trimming of U-adapter and TIR sequences, 74% (1,173,003,813 read pairs; Table 1) of the read pairs remained. Among the remaining read pairs 90% were aligned to the B73 genome containing 44,303 high-confidence gene models (Zm-B73-REFERENCE-NAM-5.0). After duplicate read removal, more than 235 million reads, i.e., read pairs and unpaired reads, remained and were used to identify germinal insertion sites at intersections of one row and one column pool. Subsequently, reads were counted in each of the 48 pools per library for insertion site identification. The MuWU analysis exclusively considered insertion sites supported by a minimum of four reads for subsequent analyses. In total, 425,924 distinct germinal insertion sites were detected in the 14 Mu*-*seq libraries tagging 36,612 (83%) of the 44,303 B73v5 genes (Table 1; Data S1). In detail, we counted insertions affecting genic regions, defined as from the start of the 5’ UTR to the end of 3′ UTR of genes, including exons and introns. Additionally, *Mu* insertions in promoter regions – up to 2,100 bp upstream of the start of the 5’ UTR – and close downstream regions of genes – up to 2,100 bp downstream of the end of the 3’ UTR – were considered. Based on the number of 425,924 insertion sites, each of the 36,612 tagged genes carried on average 12 insertional alleles (425,924 insertion sites/ 36,612 affected genes). The majority of 60% of the affected genes (21,812 of 36,612 B73v5 genes) harbored insertions in their coding sequence (Data S1). Among the 8,064 *BonnMu* F_2_-families under analysis, 98% (7,908 of 8,064 F_2_-families) carried at least one germinal insertion. Only for a minority of 2% of the *BonnMu* F_2_-families (156 of 8,064 F_2_-families), no insertion was detected. Hence on average, each *BonnMu* F_2_-family carried 53 heritable *Mu* insertions (425,924 insertion sites/ 8,064 F_2_-families). An extreme example is the *BonnMu* F_2_-family F7-4-F-1766 hosting 338 distinct *Mu* insertions in 457 different genes.

**Table 1.** Alignment statistics for Mu-seq libraries.

| Description | 14 Mu-seq libraries* |
| --- | --- |
| Raw read pairs | 1,589,204,597 |
| Read pairs after trimming | 1,173,003,813 |
| Average alignment rate | 90% |
| Read pairs and unique unpaired reads after removing duplicates | 235,240,526 |
| Number of germinal *Mu* insertions (unique insertions) | 425,924 |
| Number of *Mu*-tagged genes (unique genes) | 36,612 |

963 \* Two previously published Mu-seq libraries (Marcon *et al.*, 2020) were included in the analysis.

The genomes of the European flint lines DK105, EP1 and F7 were recently sequenced (Haberer *et al*., 2020). In light of this, we explored the number of germinal insertions sites by mapping the Mu-seq reads of the 3,456 sequenced *BonnMu* F_2_-libraries in DK105, EP1 and F7 genetic background (Table S1) to their respective genomes. Considering only genes that can be assigned to chromosomes, there are 46,726, 43,375 and 44,043 genes in the DK105, EP1 and F7 genomes, respectively. Among the 462 *BonnMu* F_2_-families in DK105 genetic background (Table S1) the MuWU analysis identified 29,986 unique insertions affecting 36% of the DK105 genes (16,704 of 46,726 genes; Data S2). Furthermore, we detected 57,806 distinct insertions in 53% of the EP1 genes (22,830 of 43,375 genes), when the Mu-seq reads of 690 *BonnMu* F_2_-families in EP1 genetic background were mapped to the EP1 genome (Data S3). Finally, the 2,304 analyzed *BonnMu* F_2_-families in F7 genetic background (Table S1) carried 64,839 unique insertions affecting 42% of the F7 genes (18,548 genes of 44,043 genes of the F7 genome; Data S4).

### *BonnMu* affected genes in flint germplasm complement the set of tagged genes identified in dent lines

Based on the analysis of 14 Mu-seq libraries in different genetic backgrounds, we uniformly mapped insertions affecting 36,612 genes to B73v5 (Table 1; Data S1). We identified varying numbers of *Mu*-tagged genes in the five different germplasm, ranging from 8,396 mutated B73v5 genes in Co125 genetic background to 32,390 B73v5 genes in B73 genetic background (Figure 1A; Data S5). This bias can be partially attributed to the fact that the Mu-seq analysis was conducted on only 576 mutagenized F_2_-families in the Co125 genetic background, whereas it was performed on 4,032 mutagenized F_2_-families in the B73 genetic background. Of the 36,612 affected genes, 5,502 (15%) were detected in *BonnMu* F_2_-families of all 14 Mu-seq libraries used in this study (Figure 1A). The number of overlapping genes significantly exceeded the expected count of 975 genes by chance (Table S2). This finding indicates the preference of *Mu* transposons for targeting specific genes across diverse inbred lines.

**Figure 1.**
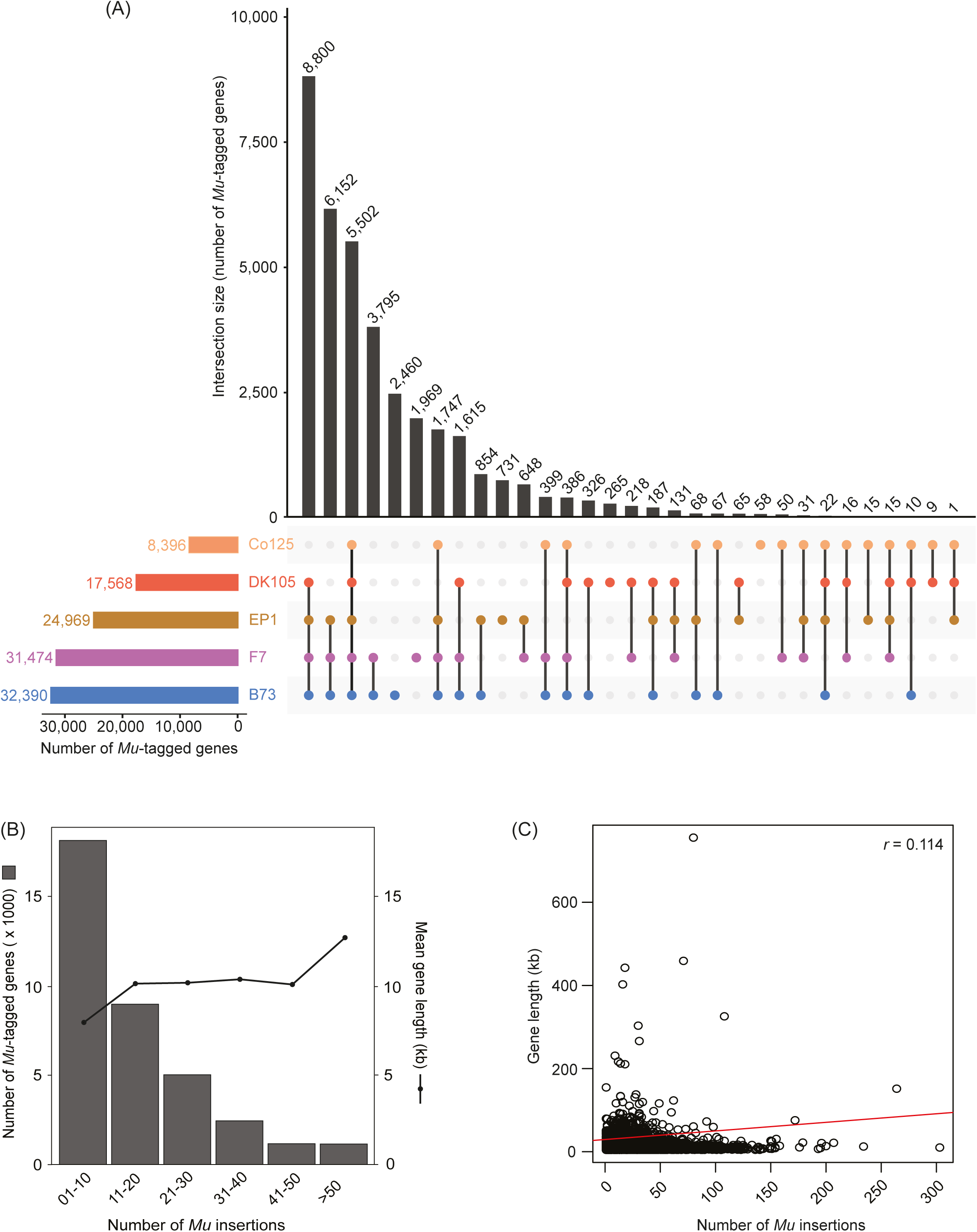
Overlap of genes affected by *Mu* insertions and distribution of insertions. **(A)** Intersections of genes, tagged in *BonnMu* F_2_-families of the two dent lines B73 and Co125 and three flint lines DK105, EP1 and F7, which have been mutagenized in this study. The UpSet plot displays 31 intersections. The lines connect overlapping genes among different genetic backgrounds. The total number of intersected genes is displayed above each bar. **(B)** Number of tagged genes and associated mean gene length plotted against the number of *Mu* insertions. **(C)** Distribution of the length of affected genes plotted against the number of individual *Mu* insertions. The calculated Pearson correlation coefficient is *r* = 0.114 (*p* < 0.001).

A higher number of 8,800 overlapping genes (24%) were identified in *BonnMu* F_2_-families of four different mutagenized germplasm: DK105, EP1, F7 and B73. A considerable proportion of *Mu*-tagged genes, that is, 16,827 (46%) of 36,612 were hit in *BonnMu* F_2_-families of Mu-seq libraries in two or three different genetic backgrounds under analysis (Figure 1A). In summary, 85% of the *Mu*-tagged genes were identified across at least two different genetic backgrounds. This result further indicated that specific genes are more prone to *Mu* transposon insertions and are affected consistently across multiple genetic backgrounds. Finally, the remaining 5,483 genes (15%) of the affected 36,612 genes were exclusively detected in one of the mutagenized inbred lines. More precisely, the majority of 2,460 genes were uniquely identified in the Mu-seq libraries in the B73 genetic background, whereas only 58 of the tagged genes were uniquely detected in the single Mu-seq library in Co125 genetic background (Figure 1A).

Among the 36,612 tagged genes, 4,027 (11%) were identified in at least one Mu-seq library of the mutagenized flint lines, i.e., DK105, EP1 and F7, but were not tagged in the set of affected genes identified in the dent lines B73 and Co125. Almost half of these genes, specifically 1,969 of 4,027 (49%), were exclusively detected in the Mu-seq libraries in the F7 background (Figure 1A). This result can be partially explained by the fact, that 2,304 *BonnMu* F_2_-families in F7 background were used for the Mu-seq experiment, whereas only 462 and 690 *BonnMu* F_2_-families in DK105 and EP1 backgrounds were analyzed, respectively (Table S1). Nevertheless, a considerable portion of 265 (7%) and 731 (18%) genes were uniquely tagged in *BonnMu* F_2_-families in DK105 and EP1 backgrounds, respectively (Figure 1A). The remaining 1,062 of the 4,027 affected genes (26%) were overlapping in *BonnMu* F_2_-families of two or three of the flint lines. Consequently, the genes affected by *Mu* insertions in flint germplasm complement the set of tagged genes identified in dent lines.

Next, we examined the distribution of all 425,924 *Mu* insertions within the 36,612 B73v5 genes, including insertions in promoter regions, i.e. within a 2,100 bp window upstream of genes and nearby downstream regions, i.e. within a 2,100 bp window downstream of genes. Among the tagged genes, nearly half of them, i.e. 49% (17,958 of 36,612) harbored 1-10 *Mu* insertions, 24% (8,908 of 36,612 genes) contained 11-insertions and the remaining 27% (9,746 of 36,612) of the genes carried at least insertions (Figure 1B). Among the latter group of genes 12% (1,160 of 9,746) carried at least 50 insertions. One extreme example is a 5,938 bp gene encoding a protein-serine/threonine phosphatase, Zm00001eb054350, carrying 303 unique insertions. Among these insertions, 38% (115 of 303 insertions) hit the coding sequence of the gene Zm00001eb054350 (Data S1).

Subsequently, we tested whether the number of *Mu* insertions was positively correlated with the length of the affected genes. To this end, we calculated the mean length of the affected genes which were grouped according to the number of insertions (Figure 1B). The five groups of genes harboring 1-50 insertions showed a comparable gene length, ranging between 7,931 bp and 10,321 bp, whereas the group of genes harboring >50 insertions exhibited an increased length of 12,595 bp. Consequently, the computed Pearson correlation across all groups of gene sizes showed a weak positive correlation between gene size and the number of *Mu* insertions (*r* = 0.114; Figure 1C).

We obtained consistent results when considering 87% of the insertions (370,396 of 425,924) affecting only genic regions, i.e., 5’ and 3’ UTRs, exons and introns (Figure S1). These insertions covered 85% of the total genes, being affected (31,126 of 36,612). In this analysis, we calculated a moderate positive correlation of *r* = 0.147 between gene size and the number of insertion sites (Figure S1).

The *BonnMu* resource provides easy-to-view photos of segregating F_2_-families at the seedling stage accessible at the genome browser at https://jbrowse.maizegdb.org/ (Marcon *et al*., 2020). Among the analyzed *BonnMu* F_2_-families various mutants were identified, such as leaf color mutants (e.g. *BonnMu*-7-C-0336) or mutants affected in shoot development (e.g. *BonnMu*-9-G-0034; Figure S2). The mutation rate in the 8,064 F_2_-families, determined by the albino and pale green leaf phenotype, was 16%. This rate aligns with previously published mutation rates (Robertson, 1983; Marcon *et al*., 2020), indicating a high transposon activity in the *BonnMu* F_2_-families.

### *Mu* insertions preferentially target the 5’ UTR of genes

Next, we investigated whether *Mu* transposons exhibit preferences for insertion sites within the maize genome. Maize has a complex genome comprising over 80% repetitive intergenic non-coding sequences (Haberer *et al*., 2020; Hufford *et al*., 2021; Chen *et al*., 2023). As a result, coding sequences constitute less than 20% of the genome. Specifically, the B73v5 genome can be subdivided into gene coding regions including 5’ UTRs (0.5%), exons (2%), introns (5%), 3’ UTRs (1%) and promoter regions (3.5%; Table S3) which are located upstream of the 5’ UTRs of genes. We further divided the promoter region into a core promoter (0.2%) and a proximal promoter (3.3%), referred to hereafter as promoterCore and promoterProx, respectively (Table S3). While the promoterCore, including the transcription start site, is located directly (1 – 100 bp) upstream of the start of the 5’ UTR, the promoterProx is 101 – 2,100 bp upstream of the 5’ UTR. Consequently, gene-coding regions and their associated promoter segments account for only 12% of the total maize genome (Figure 2A). The remaining 88% is composed of non-coding repetitive sequences (Figure 2A).

**Figure 2.**
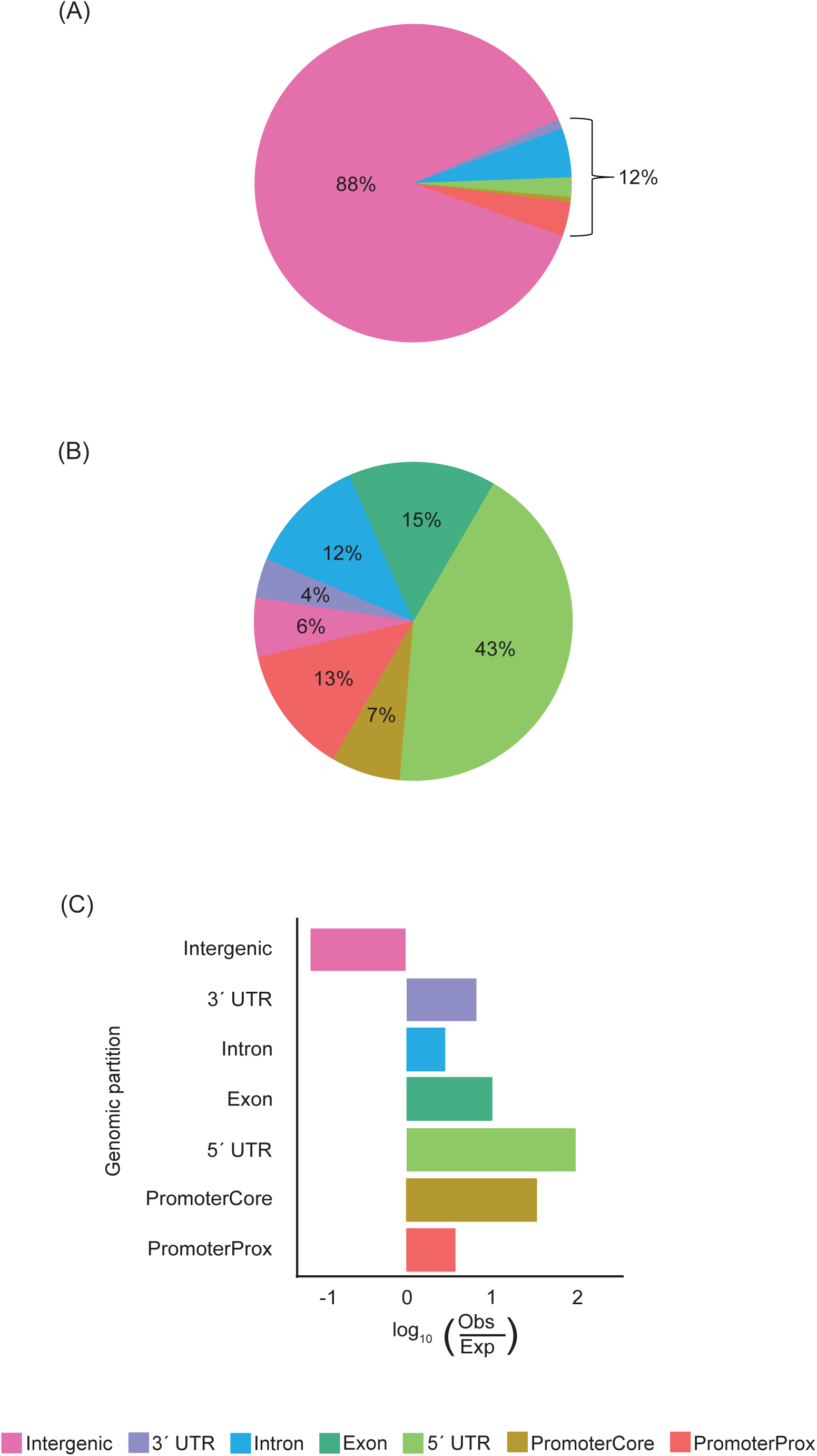
Distribution of *BonnMu* insertions across the maize genome. **(A)** Composition of the maize genome (B73v5). **(B)** *Mu* insertion sites across the genome. **(C)** Ratio of observed and expected *Mu* insertions across the genome.

To explore whether *Mu* transposons exhibit preferences for distinct categories of the coding regions or the intergenic region of the maize genome, we investigated the set of all *BonnMu* insertion sites identified in the subset of seven Mu-seq libraries consisting of 4,032 *BonnMu* F_2_-families in B73 genetic background (Data S1). Among all 774,692 somatic (not shown) and germinal insertions, a considerable proportion of 43% (331,168 insertions) affected the 5’ UTRs (Figure 2B). A comparable fraction of 15% (115,786 insertions) and 12% (95,371 insertions) tagged exons and introns of genes, while 13% (98,510 insertions) were incorporated in promoterProx sections of the maize genome. Minor fractions of insertions were identified in 3’ UTRs (4%; 27,189 insertions) and promoterCore regions (7%; 53,950 insertions). Interestingly, while 88% of the maize genome contains non-coding regions (Figure 2A; Table S3), only 6% (52,718 insertions) of the *BonnMu* insertions were detected in those regions (Figure 2B). In summary, our results indicate that intergenic insertions are underrepresented and genic regions, such as the 5’ UTRs, are frequently targeted by *BonnMu* insertions. To further support this finding and to account for the non-uniformity of intergenic *versus* genic space, we calculated the ratio between the number of observed and expected insertions per genomic partition using Pearson’s χ^2^ tests with Yates’ continuity correction. Indeed, we observed more insertions than expected for all genic and promoter partitions of the genome (Table S4; Figure 2C). For the 5’ UTRs, a number of 3,824 insertions would be expected based on the genomic composition of the maize genome (Table S4). However, we discovered significantly more than expected, specifically 331,168 insertions, exceeding the anticipated count by over 86-fold. In contrast, we expected 684,938 *BonnMu* insertion sites in intergenic regions. However, only 52,718 of such insertion sites were detected (Table S4; Figure 2C), indicating approximately 13 times fewer than expected.

### *BonnMu* insertions correspond with gene-dense telomeric regions

Previous studies described that *Mu* insertion site frequencies align with gene density (Schnable *et al*., 2009; Springer *et al*., 2018). Thus, we analyzed the distribution of *BonnMu* insertions in the B73 genetic background across all 10 chromosomes of maize by dividing each chromosome into 10k bins of 213,167 bp in size and counted the number of insertions per bin. At the heterochromatic centromeric regions of each chromosome, we predominantly detected bins containing less than 200 *BonnMu* insertions (Figure 3). In contrast, at the telomeres we frequently detected bins that carry 250 – 500 insertions (Figure 3), indicating that *Mu* elements preferentially insert into these gene-rich regions. However, there are some exceptions, i.e. bins which are in close proximity to the telomeres of the chromosomes 5-8, but lack insertions. One extreme example is the telomeric region at the short arm of chromosome 6, illustrating a 5-6 Mb window lacking insertions (Figure 3). We also observed that the distribution of *Mu* elements aligns with the chromosomal gaps in regions of chromosomes 5-8. We investigated this using the MaizeGDB JBrowse genome browser (Woodhouse *et al*., 2021) and can conclude that these gaps correspond to highly heterochromatic knob regions located at the telomeric regions of these chromosomes (Ghaffari *et al*., 2013).

**Figure 3.**
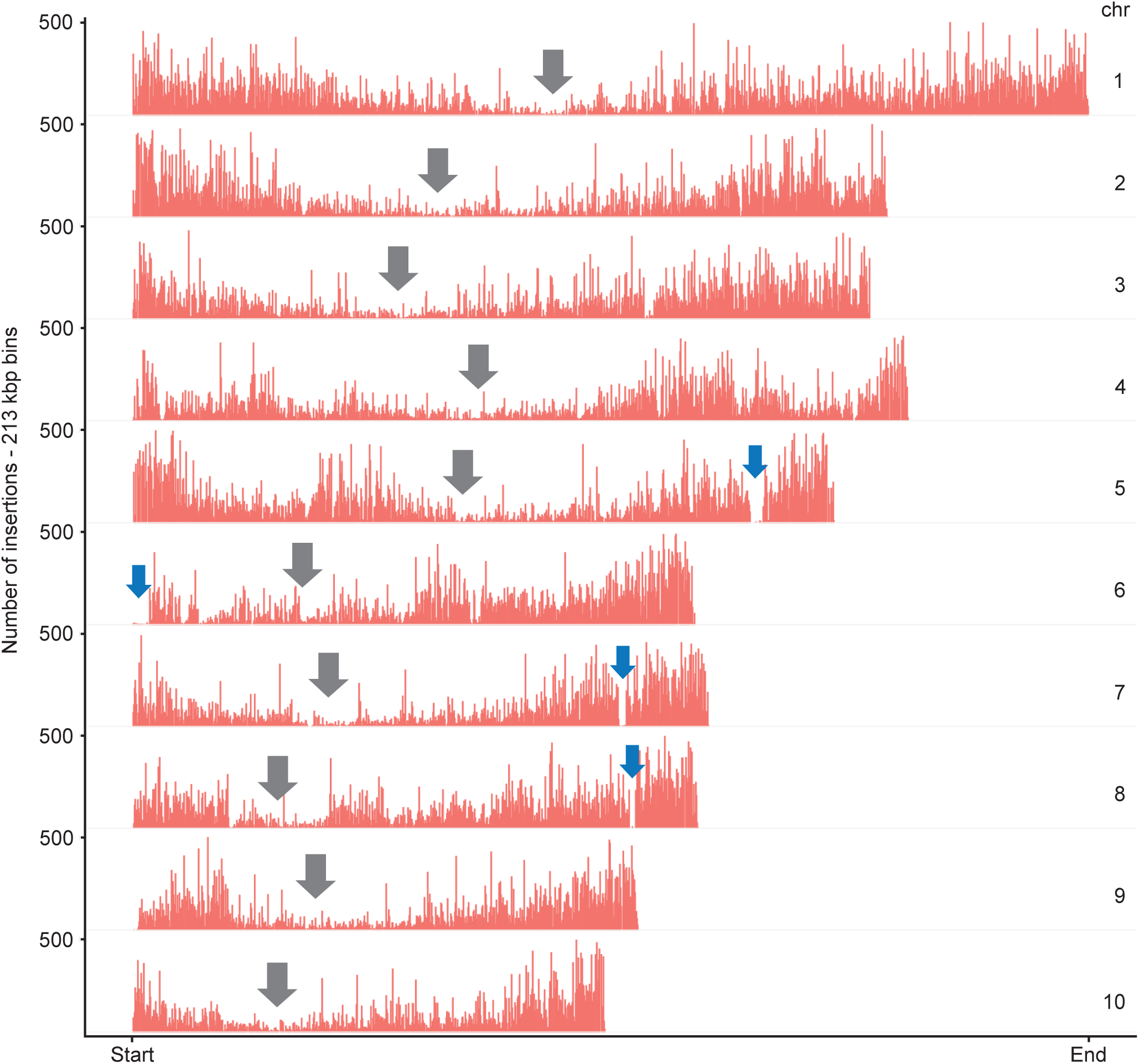
Distribution of *BonnMu* insertions across ten chromosomes. *Mu* insertions are predominantly found in the gene-rich telomer regions with fewer insertions in the heterochromatic centromere regions. Grey arrows indicate the centromeric regions in each chromosome. Blue arrows indicate gaps, i.e. regions with marginal numbers of *BonnMu* insertions, on the arms of chromosomes 5-8. These regions correspond to highly heterochromatic knob regions (Ghaffari *et al*., 2013).

### Spatial distribution of *BonnMu* insertions and chromatin accessibility across the maize genome

To gain a deeper understanding of the distribution patterns of *BonnMu* insertions within the genome, we investigated the relationship of the *BonnMu* families in B73 background to chromatin accessibility and histone modifications. To accomplish this, we utilized multiple ATAC-seq (Assay for Transposase-Accessible Chromatin using Sequencing) analyses, ChIP-seq data and published datasets related to DNA acetylation and methylation processes. ATAC-seq is used to assess open chromatin regions on a genome-wide scale (Buenrostro *et al*., 2013). In this study, we visualized the chromatin accessibility within the maize NAM (nested association mapping) population and various tissues, such as ear and leaf (Figure 4; Ricci *et al*., 2019). Additionally, the comparative analysis, juxtaposed the frequency of *Mu* insertion sites with the genome-wide distribution of chromatin modifications, including trimethylation of Lys-27 of histone H3 (Makarevitch *et al*., 2013) and histone acetylation (Zhang *et al*., 2015).

**Figure 4.**
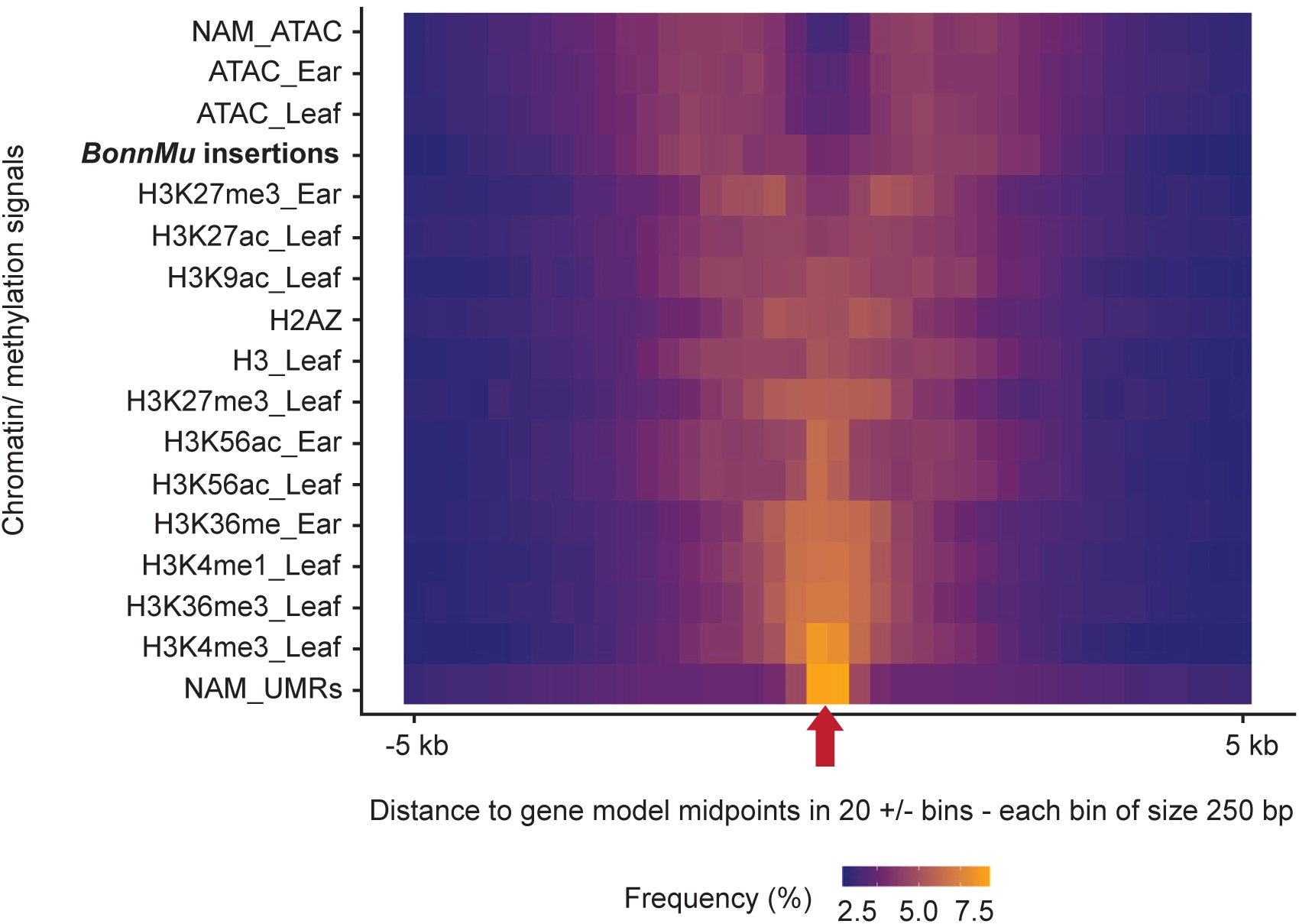
Epigenomic landscape surrounding *BonnMu* insertions in maize. Frequency distribution of *BonnMu* insertions, chromatin modifications and chromatin accessibility in relation to the entire set of genes in the maize genome. The red arrow on the horizontal center of the plot indicates the midpoint of all maize gene models. Bins of 250 bp in size are presented on both sides of this midpoint, representing a 5 kb upstream and a 5 kb downstream region of the gene midpoint. The frequency of *Mu* insertions, chromatin marks or accessibility signals is color coded (yellow = higer frequency; blue = lower frequency). NAM, nested association mapping; UMR, unmethylated regions.

In a 250 bp window around the midpoint of the maize gene models, we detected strong signals of unmethylated regions (UMRs; NAM_UMRs, Figure 4) which is in line with the high overlap of UMRs with accessible chromatin regions reported previously (Hufford *et al*., 2021). Similarly, distinct central enrichment at the gene model midpoints was observed for histone 3 modifications, such as trimethylations at Lys-4 and Lys-36 (H3K4me3_Leaf and H3K36me3_Leaf) or acetylation of Lys-56 (H3K56ac). The frequency of these histone modifications gradually diminishes in 250 bp windows both up- and downstream of the gene midpoint.

A contrasting pattern was observed for chromatin accessible signals, based on ATAC-seq datasets and *BonnMu* insertion features. While gradually increasing frequencies of ATAC-seq signals and *Mu* insertions were identified in 250 bp windows flanking the gene midpoint, there are only a few such signals and insertions in the center of the gene. Hence, *BonnMu* insertions aligned well to transposase accessible chromatin signals detected in the following datasets: ATAC_Ear, ATAC_Leaf and NAM_ATAC (Figure 4). According to Figure 4 *Mu* transposons predominantly insert at the start or end of genes, including UTRs and closely adjacent regulatory sequences. This finding is, at least in part, in line with the preference for *BonnMu* insertions targeting promoterCore and 5’ UTR regions of genes (Figure 2B). Aligning partly with the pattern observed for the *BonnMu* insertions, H3K27me3 modifications displayed tissue-specific differences in their distribution around gene midpoints. This can at least partly be explained by the reported observation that H3K27me3 seems to be less coupled to chromatin accessibility than other modification – which on average deviate from open chromatin signals only in 15-21% of cases in a tissue-specific manner in maize (Ricci *et al*., 2019).

### Mapping and validation of *Mu1* and *Mu8* transposons

The *Mu* transposon system is a powerful tool for large-scale mutagenesis in maize. Several non-autonomous *Mu* species have been comprehensively characterized (Lisch, 2015). In our study, we pinpointed the potential *Mu* species at the 425,924 distinct germinal insertion sites, tagging 36,612 B73v5 genes (Table 1; Data S1). To identify these potential *Mu* species, we associated the *Mu*-TIR sequence which was part of each Mu-seq read (i.e. flanking the gene sequence of interest), to a list of known *Mu* species (for details see “Experimental procedures”). Due to the highly conserved nature of the TIRs in all *Mu transposons* and the presence of non-specific, short TIR fragments in the Mu-seq reads, our reliable confirmation was limited to *Mu1*, *Mu8*, or *MuDR* transposons. Overall, we identified 82,285 (14.9%) *Mu1*, 1,211 (0.2%) *Mu8*, 80,347 (14.5%) *Mu8*|*MuDR* species. For the majority of 390,124 (70.4%) of the 425,924 insertion sites, we could not identify the respective *Mu* species.

For validating *BonnMu* insertions and corresponding *Mu* species, we randomly selected insertions in three distinct genes: (1) Zm00001eb052530 carrying a *Mu8* insertion in the F_2_-family *BonnMu*-2-A-0982, (2) Zm00001eb280980 and (3) Zm00001eb256020 harboring a *Mu8* and *Mu1* insertion in the mutagenized F_2_-families *BonnMu*-7-C-0459 and *BonnMu*-F7-2-F-1001, respectively. According to Data S1 following insertion identifiers, i.e. distinct 7-digit numbers, identifying unique insertions, were assigned to the three insertions: (1) *BonnMu*0031087, (2) *BonnMu*0170576 and (3) *BonnMu*0446992. The insertion identifier *BonnMu*0031087 indicates a *Mu8* insertion in the single exon of the gene Zm00001eb052530 (Figure 5A), located 157 bp downstream of the A from the ATG start codon (Data S1). A PCR-based co-segregation analysis of 11 individual plants of the segregating F_2_-family *BonnMu*-2-A-0982, identified two plants (# 1 and # 11) being homozygous for the wild type allele. Mutants were identified as heterozygotes, so both wild type and mutant specific bands were observed (# 2 - # 10; Figure 5B). Sanger sequencing (Sanger *et al*., 1977) of the *Mu*-specific PCR products confirmed that the *Mutator* insertion in this gene was caused by a *Mu8* element (Figure 5C; Data S1).

**Figure 5.**
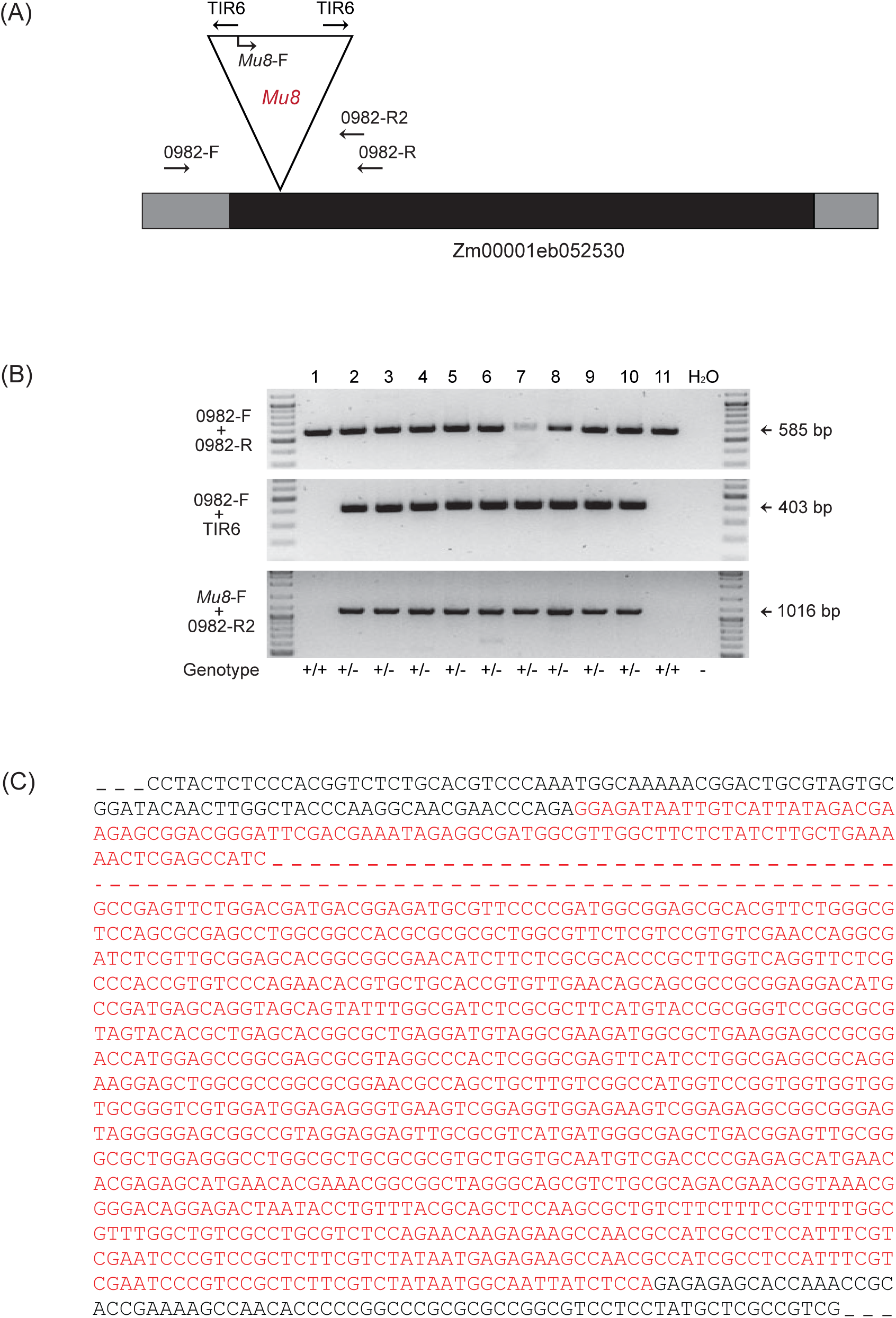
PCR-based segregation analysis. **(A)** Structure of the gene Zm00001eb052530. The single exon is illustrated as a black box and UTRs as gray boxes. The *Mu8* insertion in the exon is shown as a triangle. Gene- and TIR-specific primer sites are indicated as arrows. **(B)** PCR segregation analysis of 11 individual plants of the segregating F_2_-family *BonnMu*-2-A-0982. Gene-specific primers (0982-F + 0982-R) flanking the insertion site were combined to detect the presence of a wild type copy of the gene. Additionally, one gene-specific primer along with a TIR-specific primer (0982-F + TIR6) were used to test for the presence of an insertion in the gene. To confirm the *Mu8* insertion, a combination of a *Mu8*-specific primer and a gene-specific primer (*Mu8*-F + 0982-R2) was used. Deionized water (H_2_O) was used as a negative control. **(C)** Confirmation of the *Mu8* insertion the gene Zm00001eb052530 by Sanger sequencing. The sequence was amplified by combining two primer pairs: 0982-F + TIR6 and *Mu8*-F + 0982-R2. The sequence represents a part of the gene with the *Mu8* insertion depicted in red letters and dashed lines. The black letters represent part of the gene sequence, while dashed lines in black represent the remaining gene sequence that was not analyzed in detail.

Similarly, we confirmed the presence of another *Mu8* element in the gene Zm00001eb280980 and a *Mu1* species in the gene Zm00001eb256020 by genotyping individual plants from the F_2_-families *BonnMu*-7-C-0459 and *BonnMu*-F7-2-F-1001, respectively (Figure S3). Subsequent confirmation involved sequencing the *Mu*-specific PCR products to verify the corresponding *Mu* elements (data not shown).

## Discussion

Sequencing of 8,064 *Mu* transposon-tagged *BonnMu* F_2_-families identified 425,924 germinal insertions, representing 36,612 (83%; Table 1) of all annotated gene models of maize. This number marks a substantial increase from the prior 57% coverage (Marcon *et al*., 2020), which incorporated data from the *UniformMu* (McCarty *et al*., 2013) and the *ChinaMu* (Liang *et al*., 2019) resources. As of June 30, 2023, the *ChinaMu* database has been updated to encompass 104,294 germinal insertions, tagging 25,948 genes, constituting a coverage of 65% (http://chinamu.jaas.ac.cn). Nearly approaching whole-genome saturation, the *BonnMu* collection in maize tagged more genes than those mutagenized in rice (60%; Wang *et al*., 2013), but fewer than in Arabidopsis (>90% gene coverage; Alonso and Ecker, 2006). For rice and Arabidopsis different techniques were employed, such as the two-component transposon system Ac/Ds-based mutagenesis (van Enckevort *et al*., 2005), the Tos17 retrotransposon mutagenesis (Miyao *et al*., 2003) and the transfer-DNA insertional mutagenesis (Alonso *et al*., 2003; Toki *et al*., 2006). To track *Mu*-induced insertions in the maize genome, the *BonnMu* library uses the high-throughput sequencing strategy Mu-seq (McCarty *et al*., 2013; Liu *et al*., 2016), which has been coupled to the robust automated downstream analysis MuWU (Stöcker *et al*., 2022), accelerating the identification of germinal insertions.

A unique feature of the *BonnMu* resource is that different North American dent and European flint germplasm groups were mutagenized thereby expanding and complementing the available resource. Notably, the flint lines DK105, EP1 and F7 are adapted to the climate of central Europe (Unterseer *et al*., 2016), and they represent important founders for European breeding programs (Haberer *et al*., 2020). In this study, we demonstrated that a substantial number of 4,027 genes were exclusively tagged in at least one Mu-seq library of the mutagenized flint lines, but not in the dent lines B73 and Co125 (Figure 1A). In contrast, the mutagenized *BonnMu* F_2_-families of the dent pool contributed 2,585 *Mu*-tagged genes, which remained unaffected in any of the mutagenized flint lines. It could be worth checking for genic presence-absence variations among the genes affected by *Mu* insertions. Moderate genic presence-absence variations exist among flint and dent germplasm. Some of these genes found in either the dent or the flint pool, are expressed at high levels (Haberer *et al*., 2020), which could contribute to line-specific adaptations to environmental impacts and hence be of interest for maize improvement and breeding. Therefore, in future studies, the effect of *Mu*-tagged presence-absence genes could be investigated to identify genotype-specific mutations and their impact on the mutant phenotype.

It has been reported that *Mu* elements exhibit a pronounced preference for 5’ UTRs of genes and tend to concentrate in genomic regions with epigenetic marks of open chromatin near the transcription start site of genes (Liu *et al*., 2009; Springer *et al*., 2018; Marcon *et al*., 2020; Zhang *et al*., 2020). In support with this, we identified the vast majority of 94% of all *Mu* insertions in genic regions of the genome, while only 6% of the insertions targeted intergenic regions (Figure 2). Moreover, the distribution pattern of *BonnMu* insertions across the 10 maize chromosomes indicates that gene-dense chromosome arms are hotspots for *Mu elements*, whereas the heterochromatic centromere regions harbor fewer insertions (Figure 3). This finding is consistent with previous observations that gene-rich chromosome arms are associated with highly accumulated *Mu* elements, e.g., in the B73 (Schnable *et al*., 2009) and W22 (Springer *et al*., 2018) genomes.

Remarkably, we pinpointed chromosomal regions, specifically on chromosomes 5-8, that exhibit minimal occurrences of *BonnMu* insertions. These areas could represent highly heterochromatic non-accessible knob regions in the genome that suppress local recombination (Ghaffari *et al*., 2013). Knobs are multi-megabase tandem repeat arrays, predominantly located in mid-arm positions of chromosomes (Dawe and Hiatt, 2004). They primarily consist of two tandemly repeated DNA sequences: the 180 bp knob repeat and the 350 bp tandemly repeated element TR-1 (Ananiev *et al*., 1998). However, they have not been fully sequenced or accurately represented in genome assemblies due to the challenge of assembling their long, repetitive structures. Karyotyping has identified knob loci in maize, but there is a significant discrepancy in their representation (Ghaffari *et al*., 2013). While the physical size of knobs extends over a million base pairs, they are represented as only a few kilobases in genome assemblies. Advances in DNA sequencing technologies have significantly improved the maize B73 genome assembly, thereby reducing the initial count of over one hundred thousand gaps to just a few thousand (Schnable *et al*., 2009; Jiao *et al*., 2017; Sun *et al*., 2018). In the genome of the maize inbred line Mo17, heterochromatic knob180 and TR-1 arrays were predominantly detected in the chromosome arms of chr1L, chr4L, chr6S, chr6L, chr8L and chr9S (Chen *et al*., 2023). Similarly, the karyotype of B73 identified such tandem repeat DNA sequences, primarily on chr4L, chr5L, chr6S, chr7L, chr8L, chr9S (Ghaffari *et al*., 2013). These areas are roughly overlapping with the regions on chr5L, chr6S, chr7L and chr8L in the B73 genome (Figure 3), showing a notable absence of *BonnMu* insertions. Since knobs represent heterochromatic repeat elements, it is likely that they are not accessible for *Mu* transposon integration.

As previously shown, *Mu* elements exhibit a preference for targeting the 5’ UTRs and transcription start sites of genes (Liang *et al*., 2019; Marcon *et al*., 2020), which corresponds to open chromatin signals (Liu *et al*., 2009; Springer *et al*., 2018). To analyze the epigenomic landscape around *Mu* elements, we aligned *BonnMu* insertions with open chromatin signals and histone modifications, obtained from published datasets (Makarevitch *et al*., 2013; Zhang *et al*., 2015; Ricci *et al*., 2019; Hufford *et al*., 2021). *BonnMu* insertion patterns were in line with transposase-accessible chromatin, as demonstrated by a comparison with ATAC-seq datasets (Ricci *et al*., 2019; Figure 4). Both transposase accessible chromatin and *BonnMu* insertions showed a depletion at gene model midpoints, gradually increasing in frequency in 250 bp bins away from these midpoints. The opposite frequency was found for most of the DNA methylation marks, which were predominantly found at midpoints of gene models. This finding was previously reported by a meta-analysis of ATAC-seq signals (Ricci *et al*., 2019) and by using Micrococcal Nuclease (Rodgers-Melnick *et al*., 2016) and DNase based assays (Oka *et al*., 2017), respectively. Generally, the relationship between TEs and chromatin in maize has been shown to be markedly variable, with a complex interplay between DNA methylation, histone modifications and TEs impacting gene expression in the maize genome (West *et al*., 2014; Zhao *et al*., 2016; Noshay *et al*., 2019; Ricci *et al*., 2019).

Functional genetics experiments using mutagenized maize stocks, such as *BonnMu* F_2_-families (Marcon *et al*., 2020 and this study) or *UniformMu* stocks (McCarty *et al*., 2013) require the validation of *Mu* insertions by PCR-based genotyping (Liu *et al*., 2016). This method specifically amplifies DNA located between the highly conserved TIR sequence of the *Mu* element and adjacent regions of the genome. The TIR6 primer, which was generated based on the TIR sequences of *Mu1*, *Mu7*, *Mu3*, *Mu8* and their variants (Settles *et al*., 2004), is typically used to confirm *Mu* insertions (Figure 5; Figure S3). The TIR6 primer is mostly effective for PCR to amplify the different classes of autonomous and nonautonomous *Mu* elements (Liu *et al*., 2016). Among the classes of *Mu* elements, *Mu1*, *Mu8* and *MuDR* exhibit the highest copy numbers in the maize genome (Liu *et al*., 2009). Here, we identified *Mu1*, *Mu8* and *MuDR* elements in 30% of the *BonnMu* insertions (Data S1; Figure 5; Figure S3), facilitating future genotyping of *BonnMu* F_2_-families. For these three classes of *Mu* elements, perfect primers can be designed to validate the insertions.

In summary, the *BonnMu* resource has undergone significant expansion, providing a comprehensive assortment of Mu-seq libraries that encompass diverse genetic backgrounds. The genetic diversity represented by the different genetic backgrounds in *BonnMu* enables the identification of genotype-specific mutations in the future. The ability to tag and identify almost every gene in the maize genome is particularly noteworthy, as it enhances the resource’s utility for researchers conducting functional genomic studies.

### Experimental procedures Plant material

Mutagenized *BonnMu* F_2_-families were generated in field nurseries at the University of Bonn (Germany), Chile, Hawaii and Mexico in the years 2014-2021 as previously described (Marcon *et al*., 2020). Briefly, we obtained the F_1_-population by crossing a *Mu*-active line (*Mu^4^ per se*; Robertson, 1983) into five distinct inbred lines: B73, Co125, DK105, EP1 and F7. Then, the F_2_-population, segregating for recessive mutations, was generated by selfing all plants of the F_1_-generation. The *BonnMu* F_2_-families comprise a genetic background of 50% inbred line (e.g. B73) and 50% a *Mu*-active line.

### Construction of Mu-seq libraries

*BonnMu* libraries were constructed using the Mu-seq method (McCarty *et al*., 2013; Liu *et al*., 2016; Marcon *et al*., 2020). Briefly, we pooled a total of 6,912 *BonnMu* families in 12 Mu-seq libraries in the genetic backgrounds of B73, Co125, DK105, EP1 and F7 (Table S1). A 2-dimensional 24 x 24 grid design was utilized to pool 576 *BonnMu* F_2_-families per library (Marcon *et al*., 2020). For each Mu-seq library construction, we germinated eight seeds per family using a paper roll system (Hetz *et al*., 1996). We incubated the seedlings in a climate chamber with a photoperiod of 16 h (28 °C, 2700 lux) and a dark period of 8 h (21 °C) at 70% humidity. At 10-12 days after germination, the leaf samples were harvested and pooled based on the 24 x 24 grid design. The samples were taken from at least three seedlings of each F_2_-family. To ensure the presence of at least one mutant allele per *Mu*-tagged gene within the 3 – 8 germinated plants per F_2_-family, the probability of 99% was calculated using dbinom() and dhyper() functions in R (R Core Team, 2021; Table S5). For the precise identification of heritable germinal insertions at the intersections of rows and columns in the grid, leaf samples from independent somatic cell lineages, i.e. alternate leaves of each seedling per family, were sampled in one distinct row and one distinct column pool. By using this method, somatic insertions appeared only in a single axis of the grid and were subsequently excluded from downstream analyses. The harvested samples were frozen in liquid nitrogen and kept at -80 °C before use. For each library, the frozen leaf samples were ground manually using pre-cooled mortars and pestles.

After isolation of genomic DNA from each pool according to Nalini *et al*., (2003), the genomic DNA was randomly sheared using a Bioruptor® Pico sonication device (Diagenode) at 2s-on/2s-off setting for 2-4 cycles to obtain the fragment sizes of about 1 kb. The size of fragmented genomic DNA was analyzed by agarose gel electrophoresis after sonication. The randomly sheared genomic DNA fragments had single-stranded overhangs, which were attentively filled in using an enzyme mix (Quick Blunting™ Kit, Thermo Fisher). This process generated blunt ends, facilitating the subsequent ligation of a double-stranded universal (U) adapter.

The *Mu*-flanking amplicons were subsequently enriched through a ligation-mediated PCR (PCR-I), using a *Mu*-TIR-specific primer and a specific primer for the ligated U adapter. Then, the fragments were incorporated with a part of an Illumina sequencing adapter and a TIR sequence in the PCR-II. To minimize the number of very short *Mu*-flanking fragments, PCR-II products were purified using a CleanNGS magnetic bead-based clean-up system (CleanNA). The final PCR-III integrates the remaining sequencing adapters and 6 bp barcodes which enabled multiplexing of the 48 pools. Quality and quantity of each Mu-seq library was assessed by using a Bioanalyzer with a DNA 7500 chip (Agilent Technologies) to obtain the required concentrations for sequencing. The multiplexed Mu-seq libraries were subjected to paired-end sequencing with a read length of 150 base pairs (bp) using the HiSeq X Ten sequencing system. The raw sequencing data were stored at the Sequence Read Archive (http://www.ncbi.nlm.nih.gov/sra) under BioProject accession number PRJNA914277.

### MuWU: Identification of *Mu* insertion sites in B73, Co125, DK105, EP1 and F7

Mu-Seq reads were processed using an automated processing pipeline, referred to as Mu-Seq Workflow Utility (MuWU; Stöcker *et al*., 2022). Briefly, *Mu* insertion sites were detected based on the characteristic 9 bp target site duplications at the insertion flanking regions of *Mu* transposons. Combined with the grid design, this allowed the differentiation between germinal and somatic insertion events. After mapping all Mu-seq reads to the B73v5 reference genome (Zm-B73-REFERENCE-NAM-5.0; Zm00001eb.1; Gage *et al*., 2020), insertions were associated with specific genomic loci. In detail, we considered *Mu* insertion sites in 5’ and 3’ untranslated regions (UTRs) of genes, exons and introns, the ≤2,100 bp upstream promoter regions and ≤2,100 bp downstream regions of genes (Data S1). With the release of MuWU v1.5, we added the capability to determine the specific classes of *Mu1, Mu8* and *MuDR* elements for a detected insertion. Details of the implementation and required data are outlined in the software’s GitHub repository (https://github.com/tgstoecker/MuWU). Finally, we generated an output table of germinal insertion events, that included additional information on each event such as genomic location, associated information based on genome annotations and the most likely class of *Mu* element (Data S1).

Furthermore, we investigated germinal insertion sites by aligning Mu-seq reads from the 3,456 sequenced *BonnMu* F_2_-families in the genetic backgrounds of the flint cultivars DK105, EP1 and F7 to their corresponding genomes: DK105 (Zm00016a.1; https://download.maizegdb.org/Zm-DK105-REFERENCE-TUM-1.0/), EP1 (Zm00010a.1; https://download.maizegdb.org/Zm-EP1-REFERENCE-TUM-1.0) and F7 (Zm00011a.1; https://download.maizegdb.org/Zm-F7-REFERENCE-TUM-1.0; Haberer *et al*., 2020). The respective output tables list germinal *Mu* insertions and affected genes (DK105: Data S2; EP1: Data S3; F7: Data S4).

### Downstream analysis of *BonnMu* insertion sites

To investigate the presence of *Mu* insertions in different libraries/ genotypes, a presence/ absence intersection matrix based on GeneIDs was created. An Upset plot was generated using the UpSetR package (Conway *et al*., 2017) in R, providing a visual representation of the intersections among the 14 Mu-seq libraries and five genotypes. Correlations between the number of insertions and the length of the affected genes were calculated based on Pearson correlation coefficient (*r*) in R v4.3.1 (R Core Team, 2021). To further visualize the distribution of *Mu* insertions across various genomic partitions, i.e., exons, introns, UTRs and promoter regions, we analyzed the seven Mu-seq libraries in B73 background. To determine if the *Mu* insertion sites align with gene density, the distribution of *BonnMu* insertions in B73 background was aligned with the genes in each chromosome using the MaizeGDB JBrowse genome browser (Woodhouse *et al*., 2021). In addition, we employed the published ATAC-seq (Ricci *et al*., 2019), ChIP-seq (Makarevitch *et al*., 2013; Zhang *et al*., 2015), NAM-ATAC and NAM-UMRs datasets (Hufford *et al*., 2021) to investigate chromatin accessibility and histone modifications in relation to *BonnMu* insertions. We used the GenomicDistributions R package (Kupkova *et al*., 2022) to analyze distributions and create visualizations.

### Identification of *Mu* species per insertion

With the release of MuWU v1.5 (https://github.com/tgstoecker/MuWU) we have incorporated a new feature facilitating the identification of sub-types or specific elements within the detected *Mu* insertions. This works by supplying a set of sequences which are specific to a particular subtype/ element of the particular transposon in question (Liu *et al*., 2009). For this feature the raw input reads have to contain this sequence. However, it has to be considered that such sequences (in our usage of MuWU) are cut/ trimmed during an analysis run. Specifically, for the *BonnMu* libraries we use a 12-fold degenerate TIR primer which is trimmed away as a prerequisite before the alignment step. Therefore, based on the subtype/ element sequence association, all matching raw reads are sorted into files for the specific subtype/ element as a process uncoupled from the normal MuWU workflow. Once the insertions are identified, we associate them via their corresponding reads with all respective subtypes/ elements. Since the TIR sequence of any particular *Mu* element can vary between the left and right end of the transposon, both the “_L” (left side) and the “_R” (right side) sequence has to be considered.

While *a priori* guided determination of known *Mu* elements in our analysis is valuable to further understand the landscape of *Mu* insertions, we also support investigation of putative novel elements. Insertions which cannot be associated with a supplied species type (no match to supplied TIR sequences) are additionally investigated. Their reads are extracted and clustered and their redundancy is removed. This, in theory, allows for the detection of putative novel types or elements that were not considered by the user/ sequencing steps and can be further investigated.

### Confirmation of *Mu* insertions by PCR

We performed PCR-based confirmation of *Mu* insertions using the following *BonnMu* F_2_-families: *BonnMu*-2-A-0982, *BonnMu*-7-C-0458 and *BonnMu*-F7-2-F-1001. To this end, 12-30 seeds per segregating *BonnMu* F_2_-family were germinated using the paper roll system (Hetz *et al*., 1996). Leaf samples were harvested 10 days after germination and genomic DNA was isolated according to Nalini *et al*., (2003). Gene-specific primers flanking the *Mu* insertion sites were designed using the Primer-BLAST online tool (https://www.ncbi.nlm.nih.gov/tools/primer-blast/). To genotype the plants, three different combinations of primers were used in separate reactions: (1) gene-specific forward and reverse primer to detect the presence of the gene copy, (2) gene-specific forward and TIR6 primer and (3) gene-specific reverse and TIR6 primer, both to detect the presence of *Mu* insertions. Primer sequences are provided in Table S6. The PCR was performed using Phusion™ High-Fidelity DNA Polymerase (Thermo Fisher). The PCR products from the individual plants that showed the presence of *Mu* insertions were subjected to Sanger sequencing (Sanger *et al*., 1977). The resulting sequences were then analyzed using BioEdit software (Hall, 1999) to confirm the presence and specific locations of the *Mu* insertions

### Accession numbers

Raw sequencing data of Mu-seq libraries are stored at the Sequence Read Archive (http://www.ncbi.nlm.nih.gov/sra) with the accession number PRJNA608624.

## Acknowledgements

We thank Christa Schulz and Helmut Rehkopf (University of Bonn) for technical assistance. This work was funded by the Deutsche Forschungsgemeinschaft (DFG) grant MA8427/1-1 to CM.

## Supporting Information

The following supplementary materials can be found in the online version of this article.

**Figure S1. Number of genes affected by *Mu* insertions and distribution of insertions. (A)** Number of tagged genes and associated mean gene length plotted against the number of *Mu* insertions. Only insertions in 5’ and 3’ UTRs, exons and introns of genes were considered. **(B)** Distribution of affected gene lengths plotted against the number of individual *Mu* insertions. The calculated Pearson correlation coefficient is *r* = 0.147. The number of insertions >50 ranges from 51 to 255 *Mu* insertions per gene.

**Figure S2. Exemplary seedling mutants segregating from two *BonnMu* F_2_-families: (A)** pale green leaf mutants from *BonnMu*-7-C-0336 and **(B)** mutants affected in shoot development from *BonnMu*-9-G-0034.

**Figure S3. Confirmation of *Mu* species by PCR. (A)** Simplified gene model of Zm00001eb280980 carrying a *Mu8* element in its 5’ UTR and PCR segregation analysis of 13 segregating plants of the *BonnMu*-7-C-0459 family (lower picture). Genotyping of 13 individual plants using gene-specific primers 0459-F + 0459-R and *Mu*-specific primers 0459-F + TIR6 and 0459-R + TIR6 identified 10 plants as heterozygotes (-/+: # 1-3; # 5-6; # 8-12) and three plants as homozygous wild types (+/+: # 4; # 7; # 13). **(B)** Simplified gene model of Zm00001eb256020 tagged by a *Mu1* element in an intron. PCR segregation analysis of *BonnMu*-F7-2-F-1001 family (lower panel) identified 7 of the segregating plants as heterozygotes (+/-: # 1-3; # 7-8; # 10-11) and 5 plants as homozygous wild types (+/+: # 4-6; # 9; # 12). Exons in the upper panels of A) and B) are illustrated as black boxes and UTRs as gray boxes. The *Mu* insertions are shown as triangles. Gene- and MuTIR-specific primer sites are indicated as arrows (F/R= gene-specific forward and reverse primers).

**Data S1:** Number of *Mu* insertions and affected genes among all *BonnMu* F_2_-families in B73, Co125, DK105, EP1 and F7 genetic background. During the MuWU analysis Mu-seq reads were mapped to the genome of B73v5.

**Data S2:** Number of *Mu* insertions and affected genes among *BonnMu* F_2_-families in DK105 genetic background. During the MuWU analysis Mu-seq reads were mapped to the genome of DK105.

**Data S3:** Number of *Mu* insertions and affected genes among *BonnMu* F_2_-families in EP1 genetic background. During the MuWU analysis Mu-seq reads were mapped to the genome of EP1.

**Data S4:** Number of *Mu* insertions and affected genes among *BonnMu* F_2_-families in F7 genetic background. During the MuWU analysis Mu-seq reads were mapped to the genome of F7.

**Data S5:** Presence (1)/ absence (0) matrix to generate the Upset plot shown in Figure 1A.

**Table S1.** *BonnMu* F_2_-families used for Mu-seq library construction. Per library, 576 mutagenized F_2_-families were pooled.

**Table S2.** Number of observed and expected *Mu* tagged genes among the *Mu* insertional libraries in various inbred lines (related to Figure 1).

**Table S3.** Partition of the B73v5 genome (related to Figure 2A).

**Table S4.** Number of observed and expected *Mu* insertions across the B73v5 genome (related to Figure 2C).

**Table S5:** Calculated probability of obtaining at least one mutant allele per tagged gene among 3 – 8 germinated plants per *BonnMu* F_2_-family. The left panel, highlighted in grey, shows probabilities for scenarios with eight germinated seedlings per F_2_-family. The right panel represents probabilities when only three out of the eight seedlings per F_2_-family germinated and were subsequently harvested. WT: wild type, mut.: mutant.

**Table S6**. List of oligonucleotide primers used for PCR-based genotyping of segregating plants of three *BonnMu* F_2_-families.

## Author contributions

C.M. and F.H. established the research project. C.M., Y.N.W. and A.B. produced the Mu-seq libraries. C.M., Y.N.W. and X.D. phenotyped the *BonnMu* F_2_-families and generated seedling photos for public access via MaizeGDB.org. T.S., M.P., A.K. and H.S. conducted the bioinformatics analyses. Y.N.W. and T.S. interpreted the data and drafted the article. C.M. and F.H. contributed in data interpretation and assisted to draft the article. All authors endorsed the final draft of the article.

## Significance statement

Our resource provides a wide array of sequence-indexed mutants covering diverse genetic backgrounds, thereby facilitating functional genomics studies in maize.

