## Supplementary material for "Expanding the *BonnMu* sequence-indexed repository of transposon induced maize (*Zea mays* L.) mutations in dent and flint germplasm": Figure S1

(A)

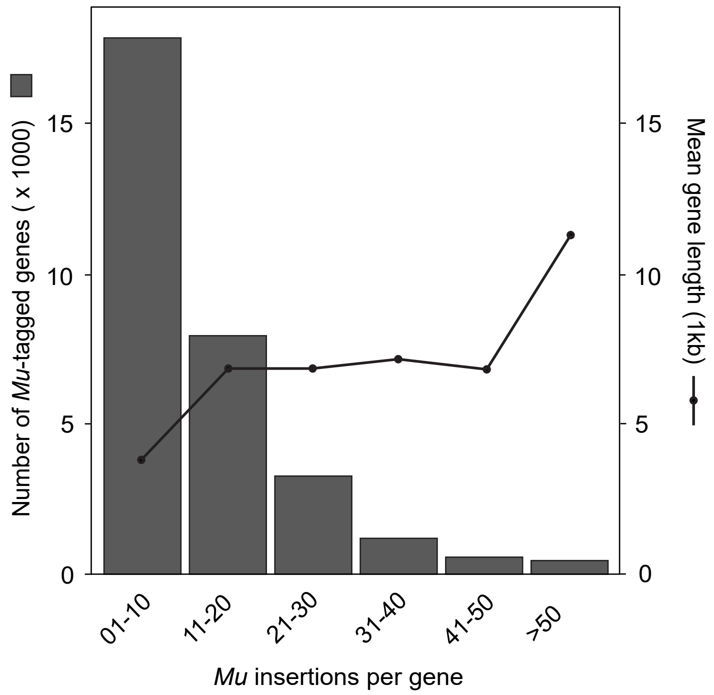


(B)

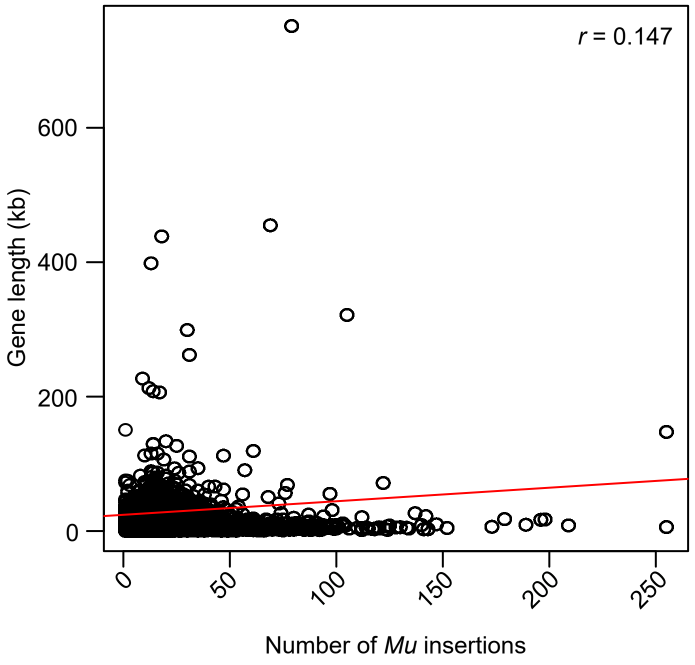


**Figure S1. Number of genes affected by *Mu* insertions and distribution of insertions. (A)** Number of tagged genes and associated mean gene length plotted against the number of *Mu* insertions. Only insertions in 5' and 3' UTRs, exons and introns of genes were considered. **(B)** Distribution of affected gene lengths plotted against the number of individual *Mu* insertions. The calculated Pearson correlation coefficient is *r* = 0.147. The number of insertions >50 ranges from 51 to 255 *Mu* insertions per gene.
