## Supplementary material for "Expanding the *BonnMu* sequence-indexed repository of transposon induced maize (*Zea mays* L.) mutations in dent and flint germplasm": Figure S2

(A)

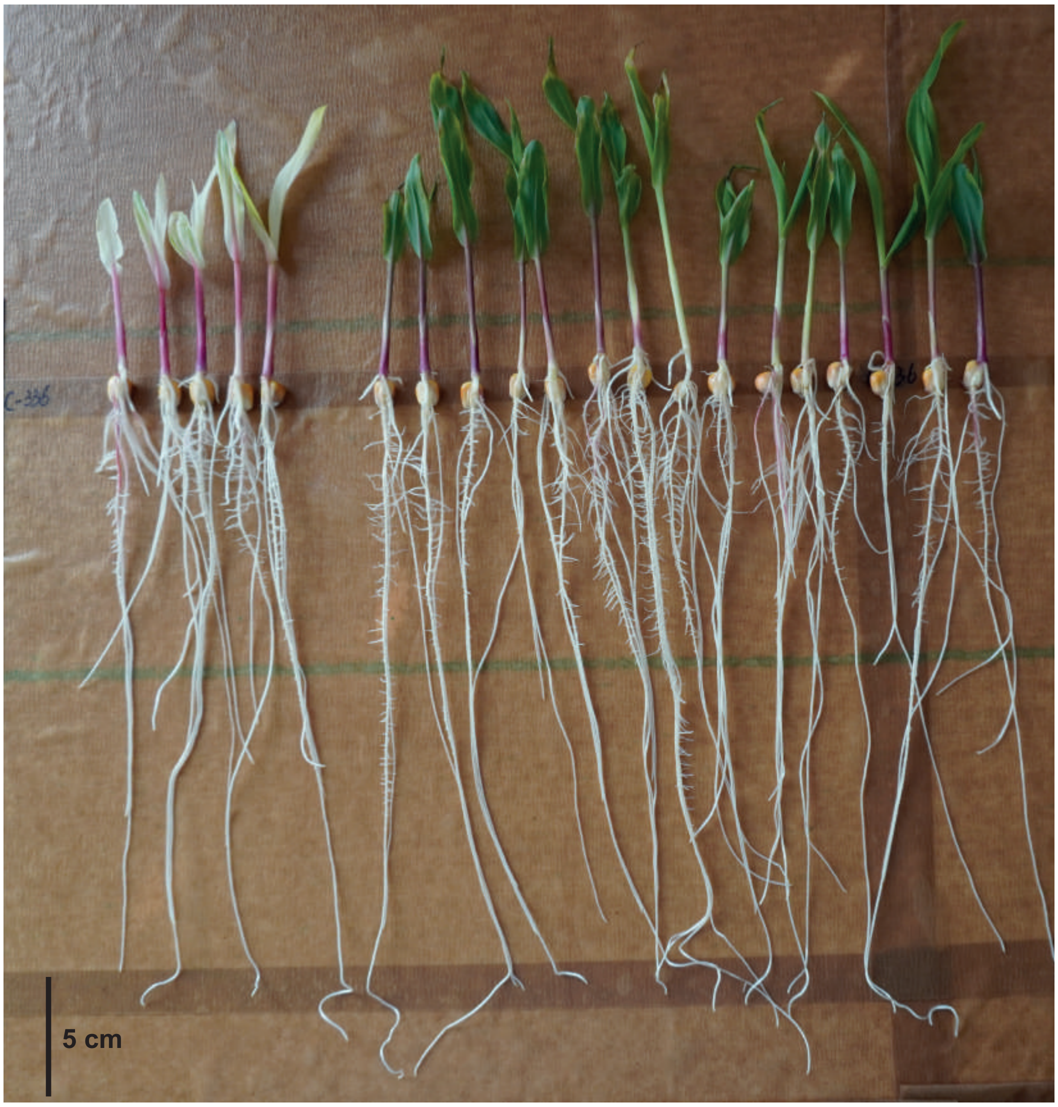


(B)

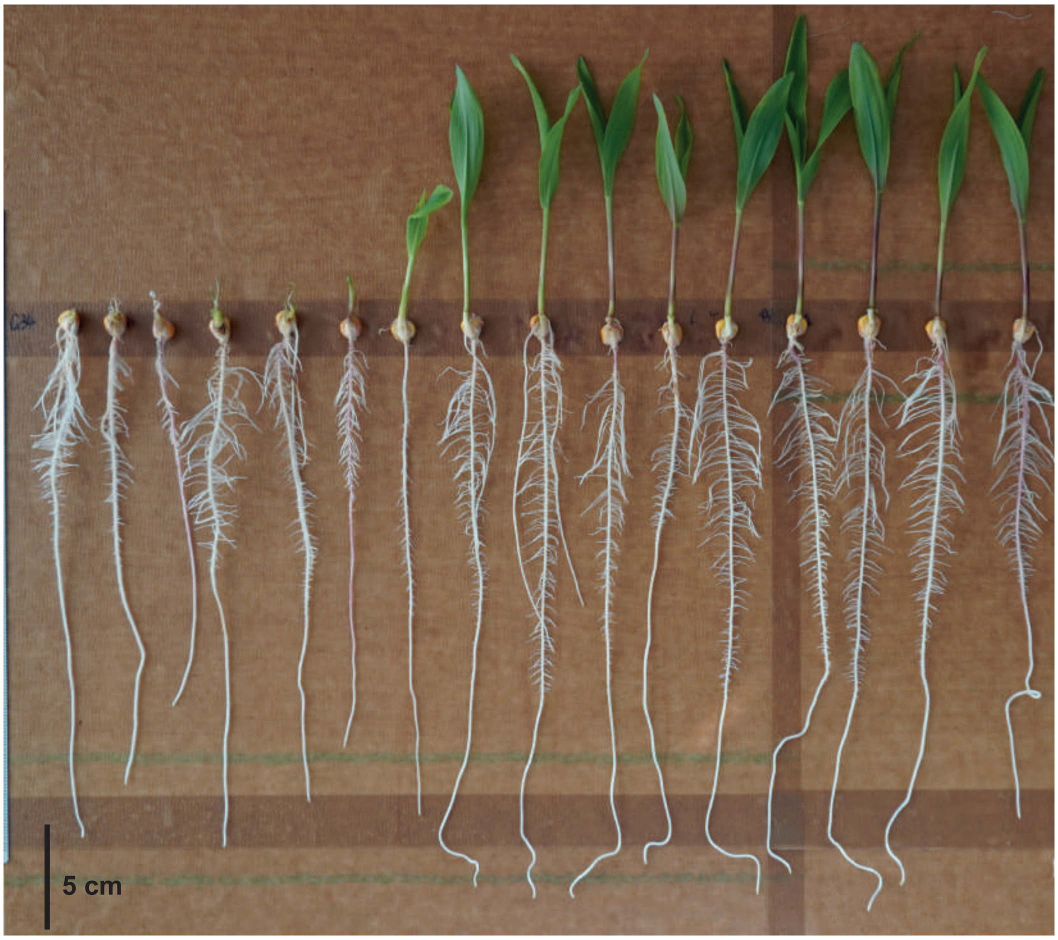


**Figure S2. Exemplary seedling mutants segregating from two *BonnMu* $F_2$-families: (A)** pale green leaf mutants from *BonnMu*-7-C-0336 and **(B)** mutants affected in shoot development from *BonnMu*-9-G-0034.
