## Supplementary material for "Expanding the *BonnMu* sequence-indexed repository of transposon induced maize (*Zea mays* L.) mutations in dent and flint germplasm": Figure S3

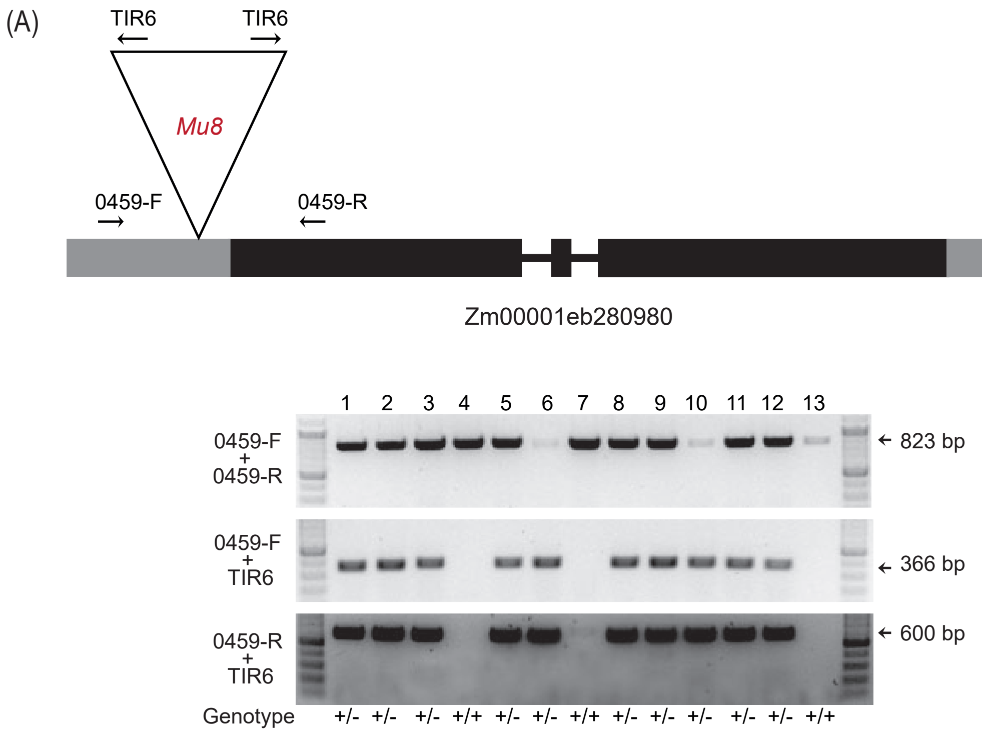


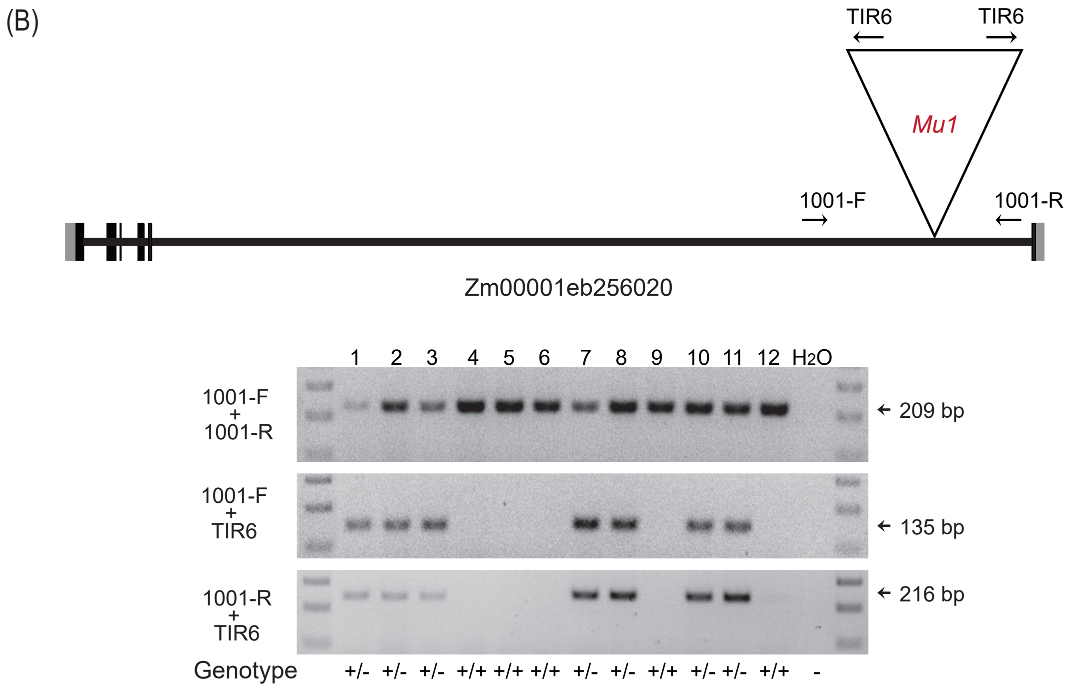


**Figure S3. Confirmation of *Mu* species by PCR**. **(A)** Simplified gene model of Zm00001eb280980 carrying a *Mu8* element in its 5' UTR and PCR segregation analysis of 13 segregating plants of the *BonnMu*-7-C-0459 family (lower picture). Genotyping of the individual plants using gene-specific primers 0459-F + 0459-R and *Mu*-specific primers 0459-F + TIR6 and 0459-R + TIR6 identified 10 plants as heterozygotes (-/+: # 1-3; # 5-6; # 8-12) and three plants as homozygous wild types (+/+: # 4; # 7; # 13). **(B)** Simplified gene model of Zm00001eb256020 tagged by a *Mu1* element in an intron. PCR segregation analysis of *BonnMu*-F7-2-F-1001 family (lower panel) identified 7 of the segregating plants as heterozygotes (+/-: # 1-3; # 7-8; # 10-11) and 5 plants as homozygous wild types (+/+: # 4-6; # 9; # 12). Exons in the upper panels of A) and B) are illustrated as black boxes and UTRs as gray boxes. The *Mu* insertions are shown as triangles. Gene- and *Mu*TIR-specific primer sites are indicated as arrows (F/R= gene-specific forward and reverse primers).
