## Supplementary material for "Expanding the *BonnMu* sequence-indexed repository of transposon induced maize (*Zea mays* L.) mutations in dent and flint germplasm": Table S1

**Table S1.** *BonnMu* F_2_-families used for Mu-seq library construction. Per library, 576 mutagenized F_2_-families were pooled.

| **Germplasm** | **Mu-seq library_ID** | ***BonnMu* F_2_-families** | **Genetic background^1^** |
| --- | --- | --- | --- |
| Dent pool | Museq_1 | *BonnMu*-1-A-0001 – *BonnMu*-1-A-0576 | (**B73** x *Mu^4^ per se*) @ |
|  | Museq_2^2^ | *BonnMu*-2-A-0577 – *BonnMu*-2-A-1152 |  |
|  | Museq_3 | *BonnMu*-3-E-0001 – *BonnMu*-3-E-0576 | (**Co125** x *Mu^4^ per se*) @ |
|  | Museq_4^2^ | *BonnMu*-4-A-1153 – *BonnMu*-4-A-1655  *&*  *BonnMu*-4-B-0001 – *BonnMu*-4-B-0073 | (**B73** x *Mu^4^ per se*) @ |
|  | Museq_5 | *BonnMu*-5-B-0074 – *BonnMu*-5-B-0650^3^ |  |
|  | Museq_6 | *BonnMu*-6-B-0651 – *BonnMu*-6-B-0922  *&*  *BonnMu*-6-C-0001 – *BonnMu*-6-C-0304 |  |
|  | Museq_7 | *BonnMu*-7-C-0305 – *BonnMu*-7-C-0805^3^  *&*  *BonnMu*-7-D-0001 – *BonnMu*-7-D-0077 |  |
|  | Museq_8 | *BonnMu*-8-D-0078 – *BonnMu*-8-D-0653 |  |
| Flint pool | Museq_DK105_EP1_1 | *BonnMu-9-G-0001* – *BonnMu-9-G-0462*  *&*  *BonnMu-9-H-0001* – *BonnMu-9-H-0114* | (**DK105** x *Mu^4^ per se*) @ |
|  |  |  | (**EP1** x *Mu^4^ per se*) @ |
|  | Museq_EP1_2 | *BonnMu-10-H-0115* – *BonnMu-10-H-0690* | (**EP1** x *Mu^4^ per se*) @ |
|  | Museq_F7_1 | *BonnMu-F7-1-F-0001* – *BonnMu-F7-1-F-0576* | (**F7**x *Mu^4^ per se*) @ |
|  | Museq_F7_2 | *BonnMu-F7-2-F-0577* – *BonnMu-F7-2-F-1152* |  |
|  | Museq_F7_3 | *BonnMu-F7-3-F-1153* – *BonnMu-F7-3-F-1728* |  |
|  | Museq_F7_4 | *BonnMu-F7-4-F-1729* – *BonnMu-F7-4-F-2304* |  |

^1^ Flint and dent lines were mutagenized by crossing with a *Mu*-active line (*Mu*^4^ per se). Selfing (@) of the F_1_-plants generated F_2_-families segregating for recessive mutations in a 3:1 ratio. Therefore, the *BonnMu* F_2_-families comprise a genetic background of 50% inbred line (e.g. B73) and 50% a *Mu*-active line. In the main text the genetic background of the different *BonnMu* F_2_-families is referred to as B73, Co125, DK105, EP1, and F7, respectively.

^2^ *Mu*-seq libraries used in a previous publication (Marcon et al., 2020).

^3^ *BonnMu* F_2_-families 5-B-0077, 7-C-0726 and 7-C-0738 failed to germinate. Consequently, they were not included in the respective Mu-seq libraries.
