## Supplementary material for "Expanding the *BonnMu* sequence-indexed repository of transposon induced maize (*Zea mays* L.) mutations in dent and flint germplasm": Table S2

**Table S2.** Number of observed and expected Mu tagged genes among the Mu insertional libraries in various inbred lines (related to Figure 1).

|  | ***BonnMu* genetic background** | | | | | **Overlap** | |
| --- | --- | --- | --- | --- | --- | --- | --- |
|  | **B73** | **Co125** | **DK105** | **EP1** | **F7** | **Expected^a^** | **Observed** |
| **Total number of *Mu* tagged genes** | 32,390 | 8,396 | 17,568 | 27,969 | 31,474 | 975* | 5,502 |

^a^ The expected overlap of *Mu*-tagged genes in five *BonnMu* genetic backgrounds was calculated using a generalized linear model with a Poisson distribution and a log-link function in R. The initial model considered each genetic background (B73, Co125, DK105, EP1, F7) as independent for the expected values, then all possible interactions were included in a second model. A χ^2^ test via the anova() function was conducted between the expected and observed numbers of *Mu*-tagged genes by comparing both models. Differences between expected and observed values were significant at α = 0.05 (* = *p* < 0.001), marked by an asterisk.
