## Supplementary material for "Expanding the *BonnMu* sequence-indexed repository of transposon induced maize (*Zea mays* L.) mutations in dent and flint germplasm": Table S3

**Table S3.** Partition of the B73v5 genome (related to Figure 2A).

| **Partition** | **%** |
| --- | --- |
| Intergenic | 88,414 |
| PromoterProx^1^ | 3,343 |
| PromoterCore^2^ | 0,178 |
| 5‘ UTR | 0,494 |
| Exon | 1,871 |
| Intron | 5,052 |
| 3‘ UTR | 0,648 |
| **Total** | **100%** |

^1^ PromoterProx: Proximal promoter region located 101 – 2,100 bp upstream of the start of the 5’ untranslated region (UTRs) of a gene.

^2^ PromoterCore: Core promoter region located 1 – 100 bp upstream of the the start of the 5’ UTR of a gene.
