## Supplementary material for "Expanding the *BonnMu* sequence-indexed repository of transposon induced maize (*Zea mays* L.) mutations in dent and flint germplasm": Table S4

**Table S4.** Number of observed and expected *Mu* insertions across the B73v5 genome (related to Figure 2C).

| **Partition** | **Observed** | **Expected** | **Log_10_** $\mathbf{(}\frac{\mathbf{Obs.}}{\mathbf{Exp.}}\mathbf{)}$ |
| --- | --- | --- | --- |
| Intergenic | 52,718 | 684,938 | -1,114 |
| PromoterProx | 98,510 | 25,899 | 0,580 |
| PromoterCore | 53,950 | 1,380 | 1,592 |
| 5‘ UTR | 331,168 | 3,824 | 1,938 |
| Exon | 115,786 | 14,491 | 0,903 |
| Intron | 95,371 | 39,141 | 0,387 |
| 3‘ UTR | 27,189 | 5,020 | 0,734 |

We used Pearson's χ^2^ test with Yates' continuity correction to calculate the ratio between the number of observed and expected insertions per genomic partition.
