## Supplementary material for "Expanding the *BonnMu* sequence-indexed repository of transposon induced maize (*Zea mays* L.) mutations in dent and flint germplasm": Table S5

**Table S****5.** Calculated probability of obtaining at least one mutant allele per tagged gene among 3 – 8 germinated plants per *BonnMu* F_2_-family. The left panel, highlighted in grey, shows probabilities for scenarios with eight germinated seedlings per F_2_-family. The right panel represents probabilities when only three out of the eight seedlings per F_2_-family germinated and were subsequently harvested. WT: wild type, mut.: mutant.

| Number of assumed mutants among eight seedlings | Probability^1^ | Probability, if 3 seedlings are harvested^2^ | | | | Probability of ≥ 1 mutant |
| --- | --- | --- | --- | --- | --- | --- |
|  |  | **3 mut.,**  **0 WT** | **2 mut.,**  **1 WT** | **1 mut.,**  **2 WT** | **0 mut.,**  **3 WT** |  |
| 0 | 0.00 | NA^3^ | NA | NA | 1.00 | 0.00 |
| 1 | 0.00 | NA | NA | 0.50 | 0.50 | 0.50 |
| 2 | 0.00 | NA | 0.10 | 0.50 | 0.40 | 0.60 |
| 3 | 0.02 | 0.02 | 0.27 | 0.54 | 0.18 | 0.83 |
| 4 | 0.09 | 0.07 | 0.43 | 0.43 | 0.07 | 0.93 |
| 5 | 0.21 | 0.18 | 0.54 | 0.27 | 0.02 | 0.98 |
| 6 | 0.31 | 0.36 | 0.54 | 0.12 | NA | 1 |
| 7 | 0.27 | 0.50 | 0.50 | NA | NA | 1 |
| 8 | 0.10 | 1.00 | NA | NA | NA | 1 |

^1^ The probability of having at least one mutant allele per tagged gene among eight seedlings was calculated using the binomial probability formula in R:

$${Bi}_{n; p}\left( k \right)=P \left( X=k \right)= \left( \begin{matrix} n \\ k \end{matrix} \right)p^{k}\left( 1-p \right)^{n-k}$$

In this formula:

n represents the number of kernels: 8.

p is the probability of each kernel harboring a mutant allele, which is 0.75.

k is the number of mutants among 8 kernels, 0-8, representing the number of mutants among the 8 kernels.

$\left( \begin{matrix} n \\ k \end{matrix} \right)$ is the binomial coefficient, representing the number of k mutants can occur in n trials.

The formula calculates the probability of having exactly k mutant alleles. To find the probability of having at least one mutant allele, we sum the probabilities for all cases where k ranges from 1 to 8

^2^ The probabilities, when only three out of eight seedlings per F_2_-family are harvest, are calculated using the following formula in R.

$$P \left( X=k \right)=\frac{\left( \begin{matrix} K \\ k \end{matrix} \right) X \left( \begin{matrix} N-K \\ n-k \end{matrix} \right)}{\left( \begin{matrix} N \\ n \end{matrix} \right)}$$

^3^ NA, combination not possible.

The final probability to have at least one mutant allele per tagged gene included among the 3-8 germinated plants per F_2_-family was calculated as:

0 x 0 + 0 x 0.50 + 0 x 0.60 + 0.02 x 0.83 + 0.09 x 0.93 + 0.21 x 0.98 + 0.31 x 1 + 0.27 x 1 + 0.1 x 1 = 0,9861 (99%).
