## Supplementary material for "Expanding the *BonnMu* sequence-indexed repository of transposon induced maize (*Zea mays* L.) mutations in dent and flint germplasm": Table S6

**Table S6.** List of oligonucleotide primers used for PCR-based genotyping of segregating plants of three *BonnMu* F_2_-families.

| **Name of primer** | **Primer sequence** | **Target sequence** | ***BonnMu* F_2_-family** | **Insertion_Identifier** | **GeneID** |
| --- | --- | --- | --- | --- | --- |
| 0982-F | GGCCTAAACTCGCAAAGGGAT | Gene-specific | *BonnMu*-2-A-0982 | *BonnMu*0031087 | Zm00001eb052530 |
| 0982-R | ATGCTTCTTGAGGCGACCAT |  |  |  |  |
| *Mu8*-F | GCCGAGTTCTGGACGATGA | *Mu8* |  |  |  |
| 0982-R2 | GACCATGGTTCTTGACGACG | Gene-specific |  |  |  |
| 0459-F | TACTACGGTTTAAGGCGTGTGG | Gene-specific | *BonnMu*-7-C-0459 | *BonnMu*0170576 | Zm00001eb280980 |
| 0459-R | AAGAGAGCAAGGGGTTTAGGC |  |  |  |  |
| 1001-F | ATATGCAGGTGAGCGGGTAG | Gene-specific | *BonnMu*-F7-2-F-1001 | *BonnMu*0446992 | Zm00001eb256020 |
| 1001-R | AAATCGAGATCGCAAGGCCA |  |  |  |  |
| TIR6 | AGAGAAGCCAACGCCAWCGCCCYATTTCGTC | *Mu*TIR |  | | |
